# Effects of fructooligosaccharides and *Saccharomyces boulardii* on the compositional structure and metabolism of gut microbiota in students

**DOI:** 10.1101/2023.12.18.572206

**Authors:** Hao Fu, Zhixian Chen, Weilin Teng, Zhi Du, Yan Zhang, Xiaoli Ye, Zaichun Yu, Yinjun Zhang, Xionge Pi

## Abstract

Fructooligosaccharides (FOS) are a common prebiotic widely used in functional foods. Meanwhile, *Saccharomyces boulardii* is a fungal probiotic frequenly used in the clinical treatment of diarrhea. Compared with single use, the combination of prebiotics and probiotics may be more effective in regulating gut microbiota. Hence, the combined use of prebiotics and probiotics has become a hot spot in synbiotics research in recent years. The present study aimed to investigate the effects of FOS and *S. boulardii—* alone or in combination—on the structure and metabolism of the gut microbiota in healthy primary and secondary school students using an *in vitro* fermentation model. The results indicated that *S. boulardii* alone could not effectively regulate the community structure and metabolism of the microbiota. However, both FOS and the combination of FOS and *S. boulardii* could effectively regulate the microbiota, significantly inhibiting the growth of *Escherichia-Shigella* and *Bacteroides*, and controlling the production of the gases including H_2_S and NH_3_. In addition, both FOS and the combination could significantly promote the growth of *Bifidobacteria* and *Lactobacillus*, lower environmental pH, and enhance several physiological functions related to synthesis and metabolism. Nevertheless, the combination had more unique benefits as it promoted the growth of *Lactobacillus*, significantly increased CO_2_ production and enhanced the functional pathways of carbon metabolism and pyruvic acid metabolism. These findings provide guidance for clinical application and a theoretical basis for the development of synbiotic preparations.

## INTRODUCTION

The importance of the intestinal microbiota has been gradually confirmed in recent years. The intestinal microbiota is composed of more than one hundred trillion microorganisms that colonize the human intestines, with a quantity equivalent to the total number of human cells (1). Hence, the intestinal microbiota constitutes an important component of the human body. Substances that cannot be degraded by the human body, such as dietary fiber, eventually reach the intestinal tract, where they are utilized and fermented by bacteria to produce substances such as short-chain fatty acids (SCFAs), vitamins, and bile acids (2). These products of bacterial metabolism have a variety of physiological effects on the host. In recent years, studies on the gut–brain axis (3), gut–liver axis (4), and gut–renal axis (5) have demonstrated that gut dysbiosis is closely related to several metabolic disorders and nervous system diseases.

The supplementation of prebiotics or probiotics is a common strategy for regulating intestinal microbiota (6). Fructo-oligosaccharides (FOS), which are non-reducing sugars composed of 3–10 fructose monomers and glucose molecules linked by β-(2,1)-glycosidic bonds, are often used as prebiotics (7). As a type of dietary fiber, FOS can enter the colon without undergoing digestion by gastric acid and endogenous enzymes and serve as a fermentation substrate for the intestinal microbiota (8). So far, FOS have shown good effects in many animal and human models. For example, it has been demonstrated that FOS can selectively promote the growth of beneficial bacteria such as *Bifidobacterium* and *Lactobacillus* in the intestinal tract (9) and also reduce the production of pro-inflammatory cytokines such as interleukin (IL)-1β, IL-6, and interferon (IFN)-γ to regulate intestinal immunity (10). In addition, FOS can also prevent oxidation and enable blood glucose homeostasis (11, 12).

*Saccharomyces boulardii* is a non-pathogenic yeast that has been widely used worldwide as a probiotic (13, 14). Previous studies have shown that *S. boulardii* can effectively alleviate various gastrointestinal diseases. The European Society of Paediatric Gastroenterology, Hepatology and Nutrition (ESPGHAN) recommends the use of *S. boulardii* in combination with other probiotics for the treatment of acute infectious diarrhea and antibiotic-associated diarrhea in children (15, 16). In addition, *S. boulardii* exerts therapeutic effects against inflammatory bowel diseases such as ulcerative colitis and Crohn’s disease (17, 18). Moreover, *S. boulardii* is often used as a feed additive for weaned piglets to improve the quality of the intestinal microbiota, increase feed conversion, and reduce diarrhea rates (19).

Although studies on FOS and *S. boulardii* have gradually become more frequent in recent years, literature on the combined use of FOS and *S. boulardii* remains rare. Although *S. boulardii* is known to have a clear preference for oligofructose (20), whether the combination of these two prebiotics has a synergistic effect warrants further investigation. In addition, most studies on the use of FOS and *S. boulardii* have been conducted in foreign populations. However, due to differences in geography, diet, age, and other factors, intestinal microbiota can differ among different populations (21). Notably, the intestinal microbiota in adolescents is unsteady and can be easily influenced by a plethora of factors. Dysbiosis of the gut microbiota can also cause diseases such as diarrhea in adolescents. Therefore, it is worth studying the specific effects of intervention with FOS and *S. boulardii* on the intestinal microbiota of adolescent populations in China. Previous studies have demonstrated that the *in vitro* fermentation model can be used to evaluate the regulatory effects of prebiotics on intestinal microbes prior to clinical trials (22). Therefore, the present study evaluated the effects of FOS and *S. boulardii* on intestinal microbes and metabolites in healthy primary and secondary school students from Hangzhou using an *in vitro* fermentation model.

## MATERIALS AND METHODS

### Materials

Live *S. boulardii* was purchased from Angel Yeast Co., Ltd. (Hubei, China); FOS, purity ≥ 95% and degree of polymerization (DP) 2–9 was obtained from Bowling Treasure Biology Co., Ltd. (Shandong, China). Tryptone, yeast extract, L-cysteine, NaCl, KH_2_PO_4_, K_2_HPO_4_, heme, MgSO_4_, CaCl_2_, and crotonic acid were all purchased from Sigma Aldrich (St. Louis, MO, USA).

### Sampling population and samples collection

Forty healthy volunteers were recruited from among primary and secondary school students in Hangzhou. All the volunteers had no gastrointestinal diseases (such as constipation or diarrhea) and had not been treated with antibiotics, prebiotics, or probiotics in the one-month preceding sample collection. The baseline characteristics of all participants are listed in Table 1. All fecal samples were quickly collected in a sterile feces sampling box, and at least 3 g of intermediate feces with little food residue and little oxygen contact was obtained during defecation. The name, gender, and age of the donor were also recorded. Samples were stored at 4° and tested within 4 h of collection. This study was approved by the Ethics Committee of Hangzhou Centers for Disease Control and Prevention (NO.202047), and written informed consent was obtained from the parents or guardians of each adolescent.

**TABLE 1.** Baseline characteristics of the study population.

| Number | Age<br>range/years | Gender | Place of residence | Gastrointesti<br>nal disorders | History of<br>antibiotics, etc. |
| --- | --- | --- | --- | --- | --- |
| 10 | 7–10 | Male | Hangzhou, China | None | None |
| 10 | 12–15 | Male | Hangzhou, China | None | None |
| 10 | 7–10 | Female | Hangzhou, China | None | None |
| 10 | 12–15 | Female | Hangzhou, China | None | None |

### Pretreatment of fecal samples

First, fresh fecal samples from student volunteers without fasting were collected and placed in two sterile centrifuge tubes. One of them was stored in the refrigerator at −80° as original fecal (OF) samples. Then, 0.8 g of fecal samples were weighed into a 10 mL sterile centrifuge tube before adding 8 mL sterile phosphate-buffered saline (PBS). The tubes were sealed and vortexed evenly. Large particles were removed by filtration using a 55mm × 28mm 300-mesh stool screen processor (Huatuo Co., Ltd. China) to obtain a 10% feces suspension. The sample pretreatment method was based on previous reports (23).

### Medium preparation

Blank control (Ctrl) medium was prepared by mixing 10 g tryptone, 2.5 g yeast extract, 1 g L-cysteine, 2 mL heme solution, 0.9 g NaCl, 0.09 g CaCl_2_·6H_2_O, 0.45 g KH_2_PO_4_, 0.45 g K_2_HPO_4_ and 0.09 g MgSO_4_·7H_2_O in 1 L of deionized water. After stirring, the culture solution was boiled and then packed into penicillin bottles using a peristaltic pump (Longer Co., Ltd., China). Continuous nitrogen was pumped into the bottles to keep an anaerobic environment during the packaging process. High-pressure steam sterilization was conducted for 15 min at 115°C and 101 kPa after sealing by a heating pressure steam sterilizer (Shenan Co., Ltd., China). The bottles were cooled and stored for later use. Meanwhile, to prepare the FOS medium, 0.8 g/100mL of FOS was added to the Ctrl medium (24).

### *In vitro* simulated fermentation by fecal microbiota

In total, 500 μL of the fecal suspension was inoculated into Ctrl and FOS culture media using disposable sterile syringes under anaerobic (Ctrl group and FOS group, respectively). A suspension of *S. boulardii* (1×10^9^ CFU/mL) was prepared, and 250 μL of the fecal suspension and 250 μL of *S. boulardii* suspension were added to the Ctrl and FOS culture media respectively (Sb group and Fsb group, respectively). Three parallel batches of each culture medium were prepared, gently shaken and mixed, and then incubated in a 37° constant temperature incubator (Biobase, China) for 24 h.

### DNA extraction and 16S rRNA sequencing of fecal and fermentation samples

After fermentation, the fermentation broth was centrifuged at 9000 r/min for 3 min (Eppendorf Centrifuge 5424, Germany), and the resultant pellet was collected for bacterial DNA extraction using the FastDNA® Spin Kit for Soil (MP Biomedicals, US) according to the manufacturer’s instructions. The V3–V4 hypervariable region of the 16S rRNA gene was amplified using the barcode fusion primers 341F (5’-CCTAYGGGRBGCASCAG-3’) and 806R (5’-GGACTACVGGGTWTCTAAT-3’). The PCR products were used for library construction and sequenced on the NovaSeq PE250 platform (Illumina, San Diego, USA) according to the protocol of Majorbio Bio-Pharm Technology Co. Ltd. (Shanghai, China). All 16S rRNA gene sequencing data was uploaded to the NCBI SRA database with the accession number PRJNA970530.

The DADA2 plug-in (25) in QIIME2 (26) was used to de-noise the optimized sequences after quality control, and amplified sequence variants (ASVs) were obtained. To minimize the effects of sequencing depth on alpha and beta diversity measure, the number of sequences from each sample was rarefied to 54000. Chao1 and Shannon index were calculated with Mothur (v1.30.1).(27) The taxonomic analysis of ASVs was performed using the Naive Bayes classifier in QIIME2 based on the SILVA database (v138) (28).

### Evaluation of SCFA content

A 2.5% (w/v) metaphosphoric acid solution was obtained by dissolving 2.5g metaphosphoric acid in 100mL deionized water. Then, 0.6464g of crotonic acid was dissolved in 100mL metaphosphate solution to obtain the crotonic acid/metaphosphate buffer solution. In total, 500μL fermentation broth was mixed with 100μL crotonic acid/metaphosphate buffer solution and then acidified in the refrigerator at −80° for 24h. After acidification, the mixture was centrifuged at 12000 r/min and 4° for 6 min, and the supernatant was filtered through a 0.22-µm microporous membrane. Finally, the filtrate was added to a gas-phase vial.

The levels of SCFAs were determined using gas chromatography (GC-2020, Shimadzu, Japan) based on the following procedure. The column temperature was maintained at 80° for 1 min and then increased to 190° at a rate of 10°/min. It was then increased further to 240° at the rate of 40°/min for 0.5 min. The FID detector and gasification chamber were maintained at 240°. The carrier gas flow rates were as follows: 20 mL/min for nitrogen, 40 mL/min for hydrogen, and 400 mL/min for air.

### Detection of fermentation gases

The fermentation bottles were removed after 24h fermentation and cooled to room temperature. The gases composition and quantity were automatically analyzed and recorded using a gas analyzer (APES-BC5-B) with five sensitive gas transduces from Empaer Technology Co., Ltd. (Shenzhen, China) according to the method described by Ye et al.(29).

### Statistical analysis

Differences in the Chao1 and Shannon index were determined using one-way ANOVA and Tukey-Kramer back-testing based on the ASV table. Structural changes in the microbial community at the genus level were analyzed, and principal coordinate Analysis (PCoA) and non-metric multidimensional scaling (NMDS) plots were obtained based on the Bray–Curtis distance algorithm (30). The abundances of all microbial phyla and genera were statistically analyzed and represented as Venn diagrams and bar plots. Linear discriminant analysis (LDA) was carried out using LEfSe software based on the ASV table, and bacteria with significant differences across different groups were identified (31). Based on Spearman correlation analysis, the correlation between each genus and SCFA and gas levels in the fermentation group were evaluated and showed in interaction network (32). All the above data analysis and mapping were carried out on the online Majorbio Cloud Platform (www.majorbio.com).

Gas and SCFA data and the results of functional prediction analysis were plotted using GraphPad Prism 8.0.1 (GraphPad Software Inc., USA) and statistically analyzed using SPSS 23.0 (IBM Corp., USA). The Shapiro–Wilk was used to test the normality of data distribution. The Fired-man test was used to test non-normally distributed data. One-way ANOVA and Tukey’s post-comparison test were used to compare normally distributed data between multiple groups. The spearman’s rank correlations between all bacterial genera, SCFAs and fermented gases were constructed into a relationship network by Cytoscape (v3.8.0) (33). A value of *P* < 0.05 was considered statistically significant.

## RESULTS

### Effects of FOS and *S. boulardii* intervention on fecal microbiota structure

The microbiota structure before and after fermentation were analyzed by 16S rRNA gene sequencing. The α-diversity analysis found the Chao1 and Shannon indexes of the Fsb group was significantly lower than that of the Ctrl group (Fig. 1A and Fig. S1), indicating that the community richness of the Fsb group was significantly reduced after fermentation. Both PCoA and NMDS plots revealed significant differences between the fecal microbiota before and after fermentation (Fig. 1B and C). Despite the high similarity between the Ctrl group and Sb groups, there were significant differences between them and other groups. At the phylum level, the fecal microbiota was mainly composed of Firmicutes, Actinobacteriota, Proteobacteria, and Bacteroidota (Fig. 1D). After *in vitro* simulated fermentation for 24 h, compared with the Ctrl group, the microbiota composition of the Sb group did not change significantly. However, the relative abundance of Proteobacteria significantly reduced in the FOS and Fsb groups, while the relative abundance of Actinobacteriota significantly increased in the FOS group, and Firmicutes significantly increased in the Fsb group (Fig. 1D). In addition, at the genus level, the number of endemic bacteria in the four fermentation groups was much lower than that in the OF group, with the number of endemic bacteria being the least in the Fsb group (Fig. 1E).

**FIG 1.**
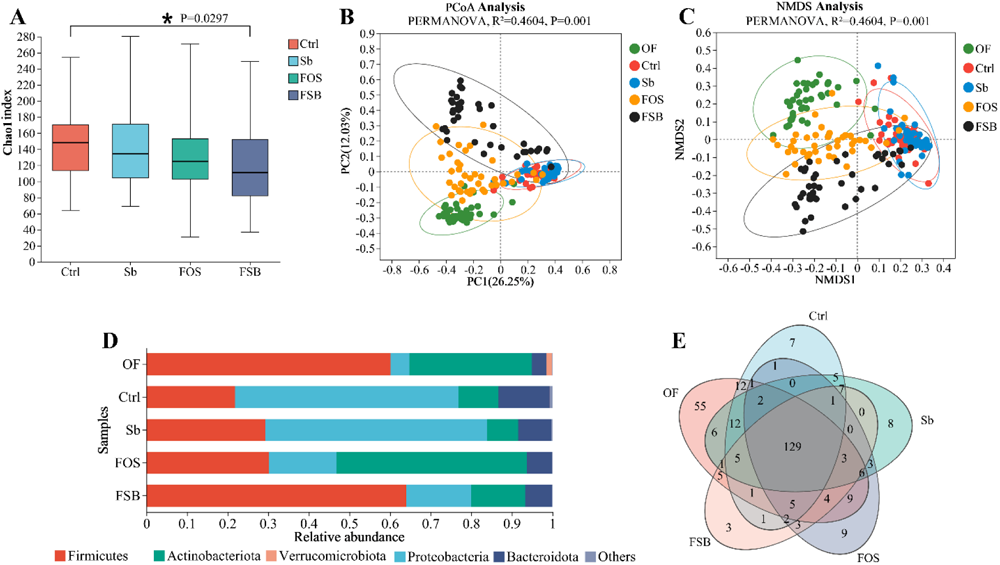
Changes in the microbial community structure before and after fermentation. (A) Chao1 index of α-diversity. ANOVA, one-way analysis of variance. Statistical significance: *, *P* ≤ 0.05. (B) β-diversity PCoA and (C) NMDS analysis. PERMANOVA, the permutational multivariate analysis of variance; (D) Composition of microbiota at the phylum level; (E) Venn diagram analysis of microbial community composition and difference at the genus level.

The genus-level microbial composition of the OF group and each fermentation group is shown in Figure 2A. In the OF microbiota, *Bifidobacterium* showed the highest relative abundance, followed by *Blautia*, *Collinsella*, *Faecalibacterium*, and others. By contrast, the relative abundance of *Escherichia-Shigella*, *Lactobacillus* and *Bacteroides* was higher in the fermentation groups. After 24 h of fermentation, significant differences in multiple bacterial genera were detected among the four fermentation groups (Fig. 2B and C). Among the genera with a relative abundance greater than 1%, *Bacteroides* showed a significantly higher relative abundance in the Ctrl group than in the other three groups (*P* < 0.001), while *Phascolarctobacterium*, *Enterococcus* and *Dorea* showed a significantly higher relative abundance in the Sb group. However, the relative abundance of *Bifidobacterium* was significantly lower in the Sb group than in the Ctrl group (*P* = 0.0081). Compared with the Ctrl group and Sb group, the FOS and Fsb groups showed a significant decrease in the relative abundance of *Escherichia-Shigella* (*P* < 0.001) and a significant increase in that of *Bifidobacterium* and *Lactobacillus*. In addition, the relative abundances of *Bifidobacterium* (*P* < 0.001), *Blautia* (*P* = 0.0072), and *Faecalibacterium* (*P* < 0.001) in the FOS group were significantly higher than those in other groups. Moreover, the relative abundance of *Lactobacillus* (*P* < 0.001) was significantly higher and that of *Collinsella* (*P* < 0.05) was significantly lower in the Fsb group than in the other groups. In general, these results suggested that the combination of FOS and *S. boulardii* can promote the reproduction of beneficial bacteria including *Bifidobacterium* and *Lactobacillus,* and play an important role in regulating the gut microbiota.

**FIG 2.**
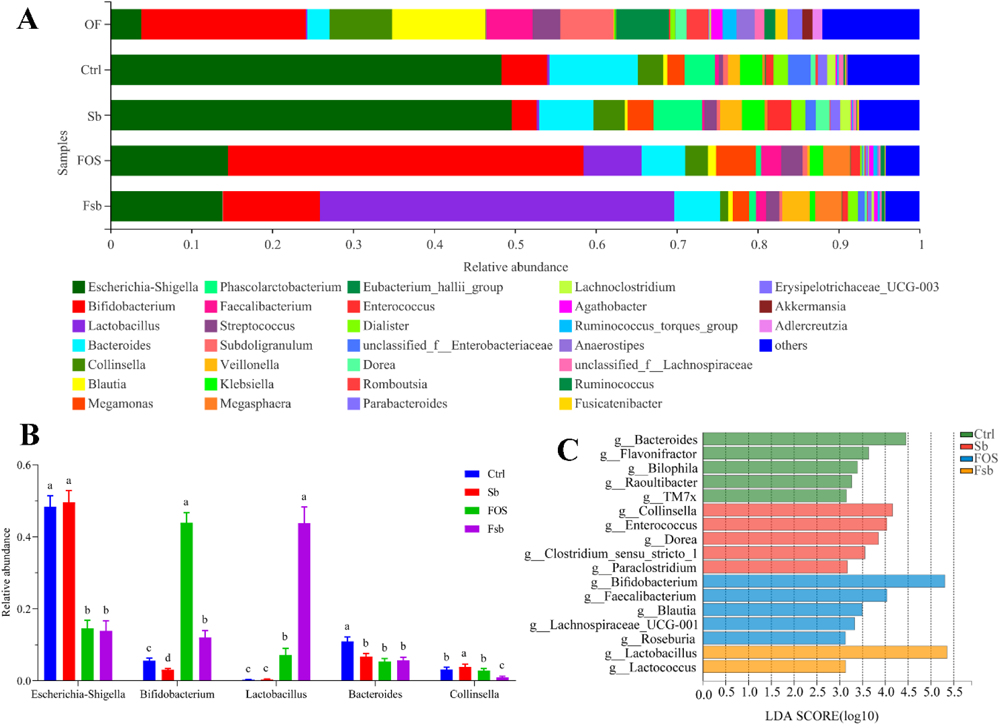
Composition of fecal microbiota and comparative analysis of multiple groups at the genus level after fermentation. (A) Composition of microbiota; (B) Multigroup comparison of five genera. Different lowercase letters represent significant differences between different groups; (C) Differential bacterial genera identified among the four groups by LEfSe analysis (LDA score > 3). LDA, linear discriminant analysis.

### Production of SCFAs by fecal microbiota during *in vitro* fermentation

The pH of the fermentation broth and the contents of six kinds of SCFAs were measured in all groups (Fig. 3). The pH values in the Sb, FOS and Fsb groups were significantly lower than those in the Ctrl group, indicating that the fecal microbiota produced more acidic products after intervention with FOS and *S. boulardii*. Furthermore, the pH value of the fermentation broth in the FOS group was significantly lower than that in the other groups (Fig. 3A). Accordingly, the FOS group showed higher levels of total SCFA production when compared with the Ctrl group (*P* = 0.0027), whereas the levels were significantly lower in the Fsb group (*P* < 0.001, Fig. 3B).

**FIG 3.**
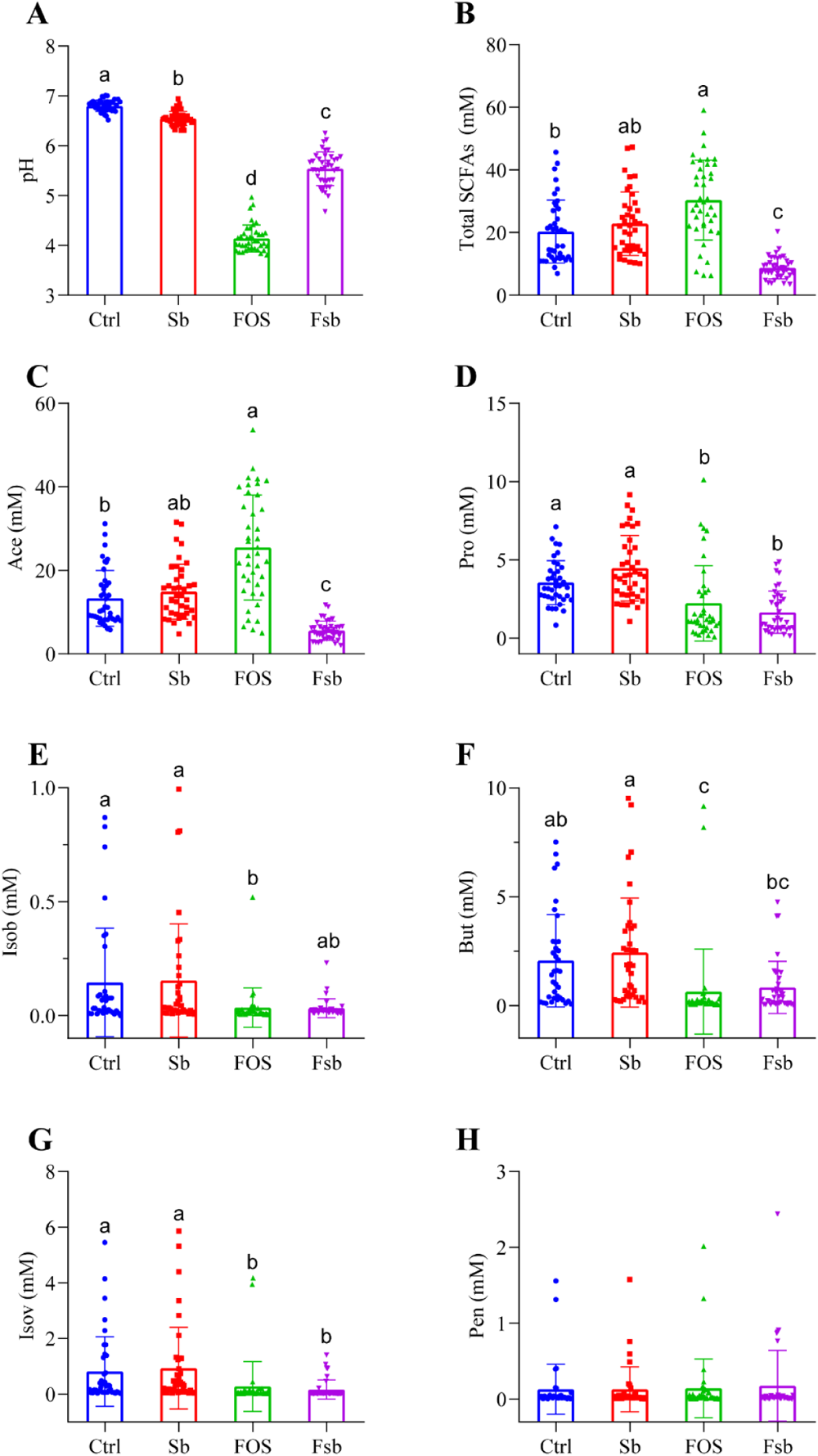
Impact of FOS and *S. boulardii* on the production of SCFAs in the *in vitro* fermentation model. (A) pH value of the fermentation broth in each group; (B) Total production of the six SCFAs in each fermentation group; The production of (C) acetic acid, (D) propionic acid, (E) isobutyric acid, (F) butyric acid, (G) isovaleric acid, and (H) pentanoic acid in each fermentation group. Different lowercase letters represent significant differences between different groups.

Among the six kinds of SCFAs, acetic acid (Ace) showed the highest production, followed by propionic acid (Pro) and butyric acid (But). Similar to total SCFA production, Ace production was significantly higher in the FOS group than in the Ctrl group (*P* < 0.001). However, Ace production in the Fsb group was significantly lower than that in the Ctrl group (*P* < 0.001, Fig.3C). Meanwhile, Pro production in the FOS (*P* = 0.0052) and Fsb groups (*P* < 0.001) was significantly lower than that in the Ctrl group (Fig. 3D). Additionally, Isob (*P* = 0.0031) and But (*P* < 0.001) production in the FOS group was also significantly lower than that in the Ctrl group (Fig. 3E-F). Moreover, the production of Isov in the FOS and Fsb groups was significantly lower than that in the Ctrl group (*P* < 0.001, Fig. 3G). There was no significant difference in the production of Pen among the four fermentation groups. Furthermore, there was no significant difference in the production of the six SCFAs between the Sb and Ctrl groups. Taken together, these results demonstrated that the combination of FOS and *S. boulardii* had different effects on the production of SCFAs by fermentation when compared with single using FOS or *S. boulardii*, especially in Ace and total SCFAs.

### Production of gases by fecal microbiota during *in vitro* fermentation

Gases are another important product of the intestinal microbiota. Hence, CO_2_, H_2_, CH_4_, H_2_S, and NH_3_ production during fermentation were also measured in Figure 4. Total gas production was significantly lower in the FOS group than in the Ctrl group (*P* = 0.0081) but significantly higher in the Sb (*P* = 0.0335) and Fsb groups (*P* < 0.001) than in the Ctrl group (Fig. 4A). As the gas with the highest production, CO_2_ accounted for the largest proportion of all gases detected. CO_2_ production in the Sb (*P* < 0.001) and Fsb groups (*P* < 0.001) was significantly higher than that in the Ctrl group (Fig. 4B). H_2_ was the second most highly produced gas. The production of H_2_ was significantly lower in the FOS (*P* < 0.001) and Fsb groups (*P* < 0.001) than in the Ctrl group (Fig. 4C). Although the production of CH_4_ was comparable among the Ctrl, Sb, and FOS groups, it was significantly increased in the Fsb group (*P* < 0.001, Fig. 4D). In addition, H_2_S and NH_3_ production in the FOS and Fsb groups was significantly lower than that in the Ctrl and Sb groups (*P* < 0.001, Fig. 4E and F). Additionally, except for the production of CO_2_, there was no significant difference in the production of the other four gases between the Sb group and the Ctrl group. Overall, these results suggested that the production of gases, such as CO_2_ and CH_4_, differed after fermenting with a combination of FOS and *S. boulardii* than after fermenting with FOS or *S. boulardii* along.

**FIG 4.**
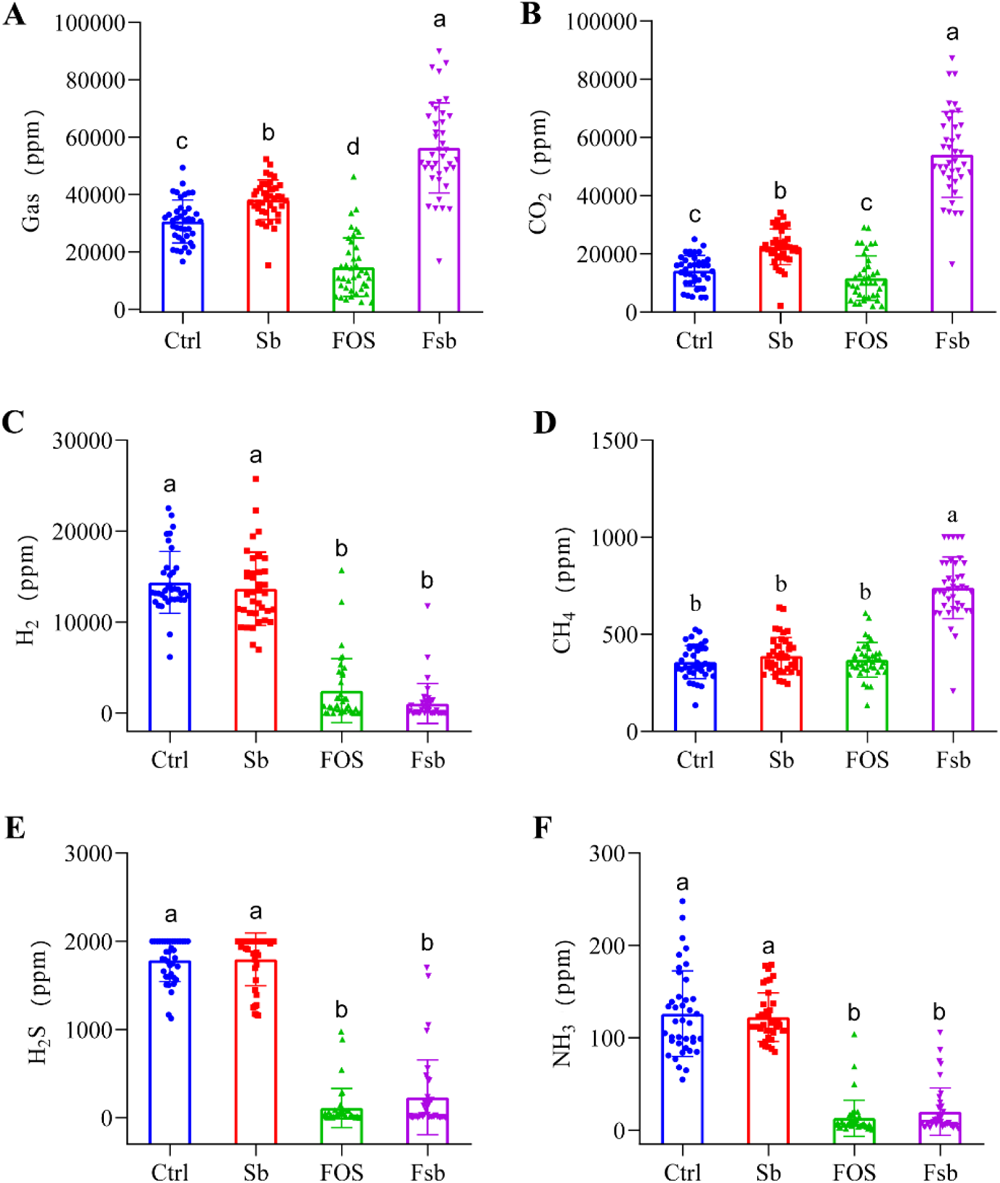
Effects of FOS and *S. boulardii* on the levels of gas produced by fecal microbiota in the *in vitro* fermentation model. (A) Total gas production in each group; Production of (B) CO_2_, (C) H_2_, (D) CH_4_, (E) H_2_S, and (F) NH_3_ in each fermentation group. Different lowercase letters represent significant differences between different groups.

### Correlation between fecal microbiota and fermentation products

We evaluated the correlation between the relative abundance of the top 15 bacterial genera and the levels of fermentation products using Spearman’s correlation analysis (Fig. 5). As important products of the intestinal microbiota, all SCFAs and gases showed a close and significant correlation with multiple bacterial genera. The well-known probiotics *Bifidobacterium* and *Lactobacillus* were negatively correlated with Pro, But, Isob, Pen, NH_3_, H_2_S, and H_2_ levels. In addition, *Bifidobacterium* was negatively correlated with Isob and CO_2_ levels and positively correlated with Ace levels. *Faecalibacterium* were also negatively correlated with NH_3_, H_2_S, CO_2_ and H_2_ Moreover, *Megamonas* and *Escherichia-Shigella*, whose relative abundance showed significant differences among the fermentation groups, were all positively correlated with But, Isob and Pen levels. Meanwhile, many genera were significantly positively correlated with H_2_S, NH_3_, CO_2_ and H_2_, including *Veillonella*, *Bacteroides*, *Escherichia-Shigella*, *Dialister*, *Phascolarctobacterium* and *unclassified_f_Enterobacteriaceae*. Further, *Megamonas* and *Klebsiella* were negatively correlated with the level of CH_4_, whereas *Collinsella* showed opposite results. These close correlations of fermentation products and bacterial genera after treatment with different combinations of FOS and *S. boulardii* demonstrated that prebiotics and postbiotics may alter the host metabolism via the action of femertation products that synthesized by the gut microbiota.

**FIG 5.**
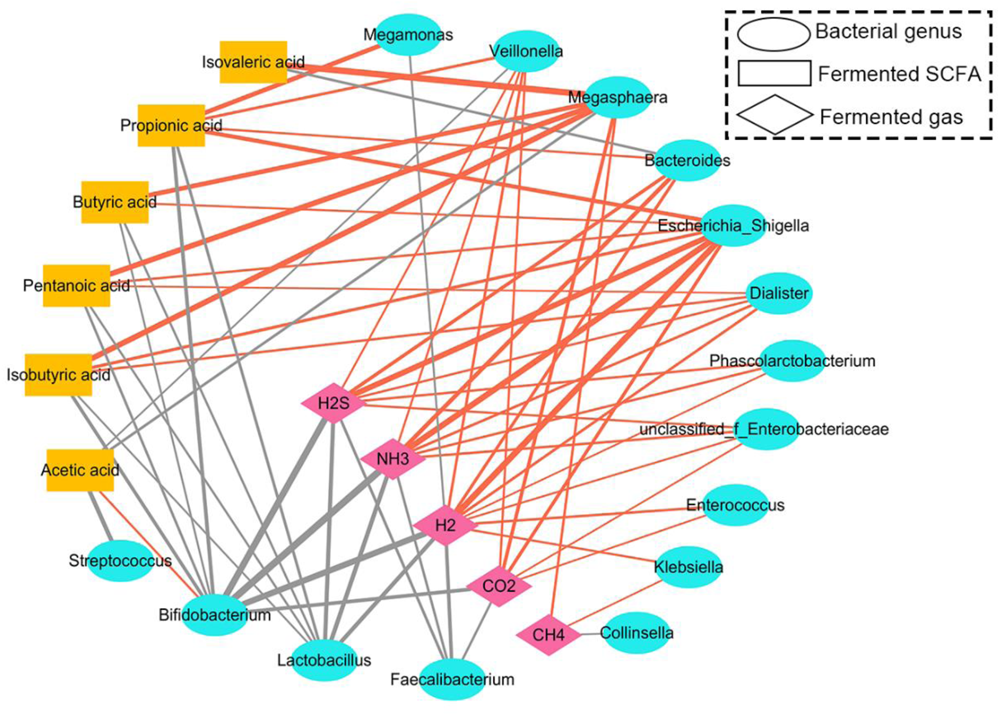
Correlation analysis of SCFAs and gases with fecal microbiota at the genus level. Spearman’s rank correlation test. Node shape indicates different genus and their fermented product. Lines between nodes represent correlations between the nodes, with line width indicating correlation magnitude, and line colors indicating positive (red) and negative (gray) correlations respectively. Only lines corresponding to correlations whose *P* value < 0.05 are drawn.

### PICRUSt2 functional prediction analysis of fecal microbiota

Based on 16S rRNA sequencing data, KEGG functions were predicted and analyzed using PICRUSt2. Functional pathways (Level 2 and Level 3) for genera with a relative abundance above 1% were compared and analyzed (Fig. S2 and Fig. 6). There were no significant differences between the functions of the Sb and Ctrl groups at Level 3 or Level 2. Only the relative abundances of metabolic pathways (*P* < 0.001) and microbial metabolism in diverse environments (*P* < 0.001) were significantly lower in the FOS and Fsb groups than in the Ctrl and Sb groups. For the FOS and Fsb groups, the relative abundances of the biosynthesis of secondary metabolites, biosynthesis of amino acids, starch and sucrose metabolism, amino sugar and nucleotide sugar metabolism, glycolysis/gluconeogenesis, purine metabolism, pyrimidine metabolism, and cysteine and methionine metabolism were significantly higher than those in the Ctrl and Sb groups (*P* < 0.05). In addition, the relative abundance of carbon metabolism (*P* = 0.0256) and pyruvate metabolism (*P* = 0.0353) in the Fsb group was significantly higher than that in the Ctrl group. The relative abundance of biosynthesis of amino acids in the FOS group was significantly higher than that in the Fsb group (*P* = 0.0256), while the relative abundance of pyruvate metabolism in the FOS group was significantly lower than that in the Ctrl group (*P* < 0.001). All these results suggested that the combination of FOS and *S. boulardii* altered different functional pathways of gut microbiota when compared with FOS or *S. boulardii* along.

**FIG 6.**
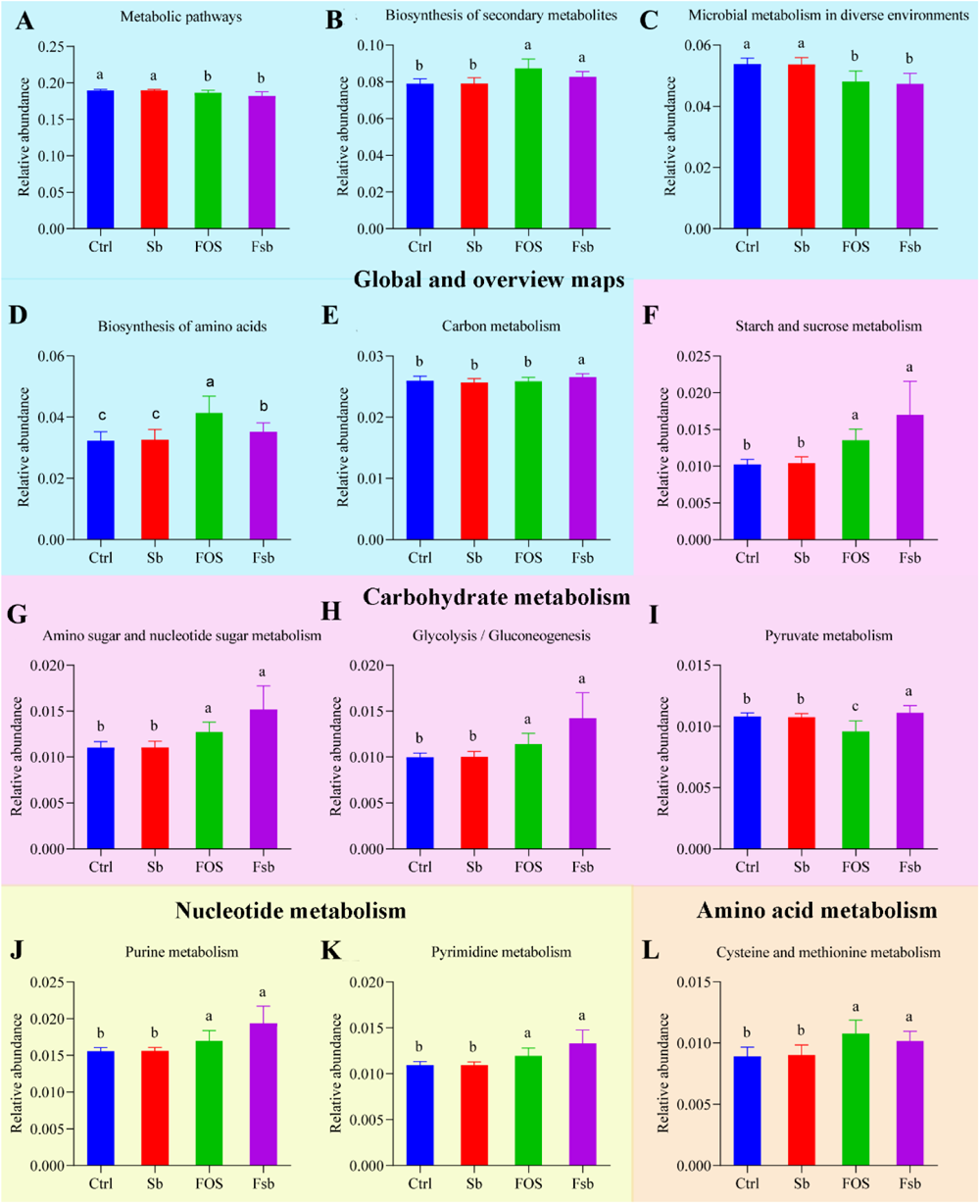
The relative abundance of KEGG functional pathway in level 3. The different background colors represent different functional pathway in level 2. Different lowercase letters represent significant differences between different groups.

## DISCUSSION

The current study explored the regulatory effect of FOS, *S. boulardii*, and their combination on the intestinal microbiota of healthy primary and secondary school students using an *in vitro* fermentation model. The results showed that the community structure of the fecal microbiota can be significantly altered during fermentation by intervention with FOS, *S. boulardii*, and their combination, which leading to changes in fermentation products and functional metabolic pathways.

The addition of FOS or *S. boulardii* alone significantly altered the composition and diversity of the fermented fecal microbiota compared to their combination. On the one hand, as a widely used and studied prebiotic, FOS has been proven to promote the growth of *Bifidobacterium* and *Lactobacillus*, which are recognized as important probiotics (34–36). The experimental results of this study also corroborated this finding. In addition, the relative abundance of *Faecalibacterium* and *Blautia* increased significantly in the FOS group. *Faecalibacterium*, an anti-inflammatory symbiotic bacterium located in the human intestines (37), is an important producer of butyric acid (38, 39). *Blautia* can produce acetic acid in the human intestine via microbial fermentation and promote the absorption of nutrients (40), (41). These SCFAs are crucial linkages between the host and gut microbiota because they are the primary degradation product of carbohydrates (42). They also play an important role in enhancing the barrier integrity of the gut epithelium and occasionally increasing the systemic IgG or IgA to regulate pathogen-specific immune reponses (43, 44). In addition to promoting the growth of beneficial bacteria, FOS also significantly reduced the relative abundance of *Escherichia-Shigella*, a common conditional pathogen (45) that can destroy intestinal epithelial tissue by promoting inflammation and subsequently cause diarrhea and other diseases (46). On the other hand, *S. boulardii*, as a fungal probiotic with therapeutic effects, is widely used in the adjuvant treatment of various intestinal diseases, such as antibiotic-associated diarrhea and gastrointestinal disorders (47, 48). It is well known that the gut microbiome of adolescents and children is unstable, and various intestinal diseases are often caused by gut microbiota dysbiosis (49). However, in our *in vitro* fermentation model, the addition of *S. boulardii* alone did not provide much benefit in regulating the fecal microbiota among healthy students. The relative abundance of *Bifidobacterium* decreased significantly in the Sb group, while that of *Collinsella* and *Dorea* increased significantly. *Collinsella* was reported associated with poor metabolic status, elevated cholesterol and low-density lipoprotein levels, and cardiovascular diseases (50). *Dorea* has been reported to show significant elevations in the intestines of adolescent patients with irritable bowel syndrome, and its excessive proliferation can be deleterious to intestinal health (51). One of the reasons for these results could be that the initial culture medium contained only nitrogen sources, whereas only a small amount of carbon sources present in the fecal diluent. This may have affected the proliferation of *S. boulardii* and the effectiveness of this probiotic. Fermentation results from the Fsb group seemed to validate this hypothesis.

Fortunately, the combination of FOS and *S. boulardii* indicated significantly different results when compared with the other groups. The relative abundance of *Escherichia-Shigella* was significantly decreased, *Bifidobacterium* and *Lactobacillus* were distinctly promoted in the fecal microbiota of healthy students. Interestingly, earlier studies have reported the interaction between yeast and *Lactobacillus* in some fermented foods (52, 53). For example, *S. boulardii* can promote the growth of *Lactobacillus* during milk fermentation (54), and yeast can provide the CO_2_ and amino acids necessary for the growth of *Lactobacillus*(55, 56). In the current study, the Fsb group produced much more CO_2_ than the other groups. An acidic environment with a pH of 4.5–6.5 is more conducive to the growth of *Bifidobacterium* and *Lactobacillus*(57), and also fitted to the pH of the human intestine ranges from 5 to 7 (58). The fermentation broth of the Fsb group was more effective in maintaining the pH within this range, while that of the FOS group was more acidic. In addition, unlike *S. boulardii* alone, the combination of FOS and *S. boulardii* significantly reduced the relative abundance of *Collinsella*, which demonstrated powerful inhibition of potentially harmful bacteria. In summary, the yeast *S. boulardii* combined with FOS showed probiotic characteristics in regulating the gut microbiota.

The intestinal microbiota affects human health by digesting dietary fiber and other substances in unabsorbed food residue to produce SCFAs and gas, which are essential metabolites for the human body (59). Therefore, the composition of these metabolites were mediated by gut microbiota and were closely related to physiological functions of body. Ace is among the most important SCFAs in the human cecum. It was used as a substrate for the synthesis of hepatic glycogen, long-chain fatty acids, glutamine, and glutamic acid, and even could trigger an immune response against potentially harmful bacteria (60–63). The FOS group produced a large amount of Ace, which significantly increased the total production of SCFAs. The interaction network of gut microbiome and microbial metabolites also revealed the positive association between *Bifidobacterium* and Ace. Meanwhile, the relative abundance of *Bifidobacterium* was significant increased in the FOS group. *Bifidobacterium* can metabolize carbohydrates to produce acetic acid and lactic acid (64), which subsequently decreases the environmental pH and further promotes the growth of *Bifidobacterium*. The excessive growth of *Bifidobacterium* and the decrease in the pH of the fermentation broth, which suppressed the growth of other intestinal bacteria and led to a significant decrease in the production of several other SCFAs. Although the pH of the Fsb group was also significantly lower than that of the Ctrl and Sb groups, the contents of the six SCFAs did not increased in the Fsb group compared to these two groups. We speculated that the Fsb group generated a large amount of CO_2_ during the fermentation, which was confirmed in Figure 4B. The increased pressure within the closed fermentation environment may have led to the partial dissolution of CO_2_ and eventually decreased the pH of the fermentation broth. Moreover, because the relative abundance of *Lactobacillus* was significantly increased in the Fsb group, the fermentation broth of the Fsb group may have contained high levels of lactic acid. Lactic acid is an important intermediate of systemic metabolism (65), and it can be metabolized into other organic acids by intestinal microbes (66). However, the proliferation of *Lactobacillus* may have inevitably reduced the area available for colonization by other acid production bacteria and eventually led to the decrease in the content of the other five SCFAs, except pentanoic acid.

As another important metabolic product of the intestinal microbiota, gases can have various physiological, pathogenic, or therapeutic effects based on their type, concentration, and volume (67). In the simulated *in vitro* fermentation model, the addition of *S. boulardii* led to more CO_2_ production, especially in the presence of FOS. As an inert gas, CO_2_ is a key molecule in many metabolic processes (68). and it can promote intestinal peristalsis through the volume effect. Excess CO_2_ can be quickly eliminated by the body (67). and typically does not harm the human body. Meanwhile, CO_2_ is beneficial to gut microbiota homeostasis during colonoscopy in healthy subjects, which improves the relative abundance of anaerobic probiotic such as *Bacteroides caccae* and *Bacteroides thetaiotaomicron* (69). However, methanogens in the intestine can use CO_2_, H_2_, and Ace to produce CH_4_ (70, 71). This could explain why the production of H_2_ and acetic acid decreased significantly in the Fsb group, while that of CH_4_ increased significantly. Previous studies have shown that CH_4_ has anti-inflammatory, anti-oxidative, and anti-apoptotic effects (72). Moreover, bacteria can accelerate the production of ATP through CH_4_ production and improve the energy-harvesting efficiency of the host (73). Furthermore, the production of H_2_S and NH_3_ decreased significantly in the FOS and Fsb groups. Current evidence reveals that the H_2_S and NH_3_ produced by intestinal bacteria are related to many gastrointestinal diseases such as ulcerative colitis, Crohn’s disease, and irritable bowel syndrome. Excessive H_2_S and NH_3_ can destroy the epithelial structure of the colon and increase intestinal permeability, increasing susceptibility to microbial infections and colon cancer (74, 75). Hence, these findings demonstrated that the combination of FOS and *S. boulardii* can improve intestinal health by reducing harmful gases.

In addition to SCFAs and gas, we also preliminarily explored the other metabolic functions of the fecal microbiota in the fermentation system based on KEGG functional prediction analysis. The relative abundance of functional pathways involved in amino acid, saccharide, and nucleotide synthesis and metabolism increased significantly in the FOS and Fsb groups, indicating that FOS and the combination of FOS and *S. boulardii* could promote microbial synthesis and metabolic function. However, the combination of FOS and *S. boulardii* also increased the relative abundance of functional pathways related to carbon metabolism and pyruvate metabolism, while FOS alone significantly decreased the relative abundance of functional pathways related to pyruvate metabolism. Hence, it suggests that the Fsb group with large number of *Lactobacillus* enriched may convert pyruvate into lactic acid under anaerobic conditions.

Overall, FOS, as a common prebiotic, can inhibit the growth of the harmful bacteria *Escherichia-Shigella* and the excessive production of the harmful gases H_2_S and NH_3_. Meanwhile, it can also promote the growth of the beneficial bacteria *Bifidobacterium*, *Lactobacillus*, *Faecalibacterium*, and *Blautia* and the production of acetic acid. As a fungal probiotic, *S. boulardii* alone could not effectively regulate the structure and metabolism of the fecal microbiota in the absence of a carbon source. However, when enough FOS and *S. boulardii* were added together to the *in vitro* fermentation system, in addition to the beneficial effects of FOS itself, the combination of FOS and *S. boulardii* could significantly promote the growth of *Lactobacillus* and optimize the environmental pH, further enhancing the relative abundance of functional metabolic pathways. However, the combination of FOS and *S. boulardii* reduced the content of SCFAs and acidic metabolites produced by the fecal microbiota, which warrants further investigation. Additionally, further research is also needed to understand how FOS and *S. boulardii* should be mixed to achieve the optimal regulation of gut microbiota. Nevertheless, these results provide a theoretical basis for the development of *S. boulardii* and FOS as synbiotics and provide a strategy for further improving intestinal microbiota in teenagers.

## Conclusions

In a simulated *in vitro* intestinal fermentation system based on nitrogen sources, FOS alone and the combination of *S. boulardii* and FOS could reduce the environmental pH, significantly inhibit the growth of the harmful bacteria *Escherichia-Shigella* and *Bacteroides* and the excessive production of the harmful gases H_2_S and NH_3_, and significantly promote the growth of the beneficial bacteria *Bifidobacterium* and *Lactobacillus*. While FOS could promote the growth of *Bifidobacterium* and increase the relative abundance of the beneficial bacteria *Faecalibacterium* and *Blautia* and the production of Ace, the combination of *S. boulardii* and FOS could also promote the growth of *Lactobacillus* and significantly inhibit the growth of the harmful bacteria *Collinsella*. Hence, the combination could further promote metabolic functions while inhibiting the production of SCFAs. However, *S. boulardii* alone could not effectively regulate the structure and metabolism of the intestinal flora. In conclusion, *S. boulardii* and FOS could be applied as symbiotics to regulate the intestinal microbiota of healthy adolescents in China, but the optimal concentration of their combination and related physiological functions require further study.

## ACKNOWLEDGMENTS

The authors would like to thank Shanghai Majorbio Bio-pharm Technology Co.,Ltd for DNA sequencing services.

## AUTHOR ORCIDs

Hao Fu http://orcid.org/0000-0001-7576-0942 Xionge Pi http://orcid.org/0000-0003-1120-0465

## AUTHORS CONTRIBUTIONS

Hao Fu, Data curation, Methodology, Data analysis, Writing – original draft, Writing – review and editing | Zhixian Chen, Data curation | Weilin Teng, Project administration, Samples collection | Zhi Du, Investigation, Validation | Yan Zhang, Resources, Supervision | Xiaoli Ye, Methodology, Resources | Zaichun Yu, Methodology, Validation | Yinjun Zhang, Methodology, Resources | Xionge Pi, Conceptualization, Funding acquisition, Supervision, Validation, Writing – review and editing.

## DATA AVAILABILITY

All 16S rRNA gene sequencing data was uploaded to the NCBI SRA database with the accession number PRJNA970530.

## ETHICS APPROVAL

This research was approved by the Ethics Committee of Hangzhou Center for Disease Control and Prevention (No. 20047).

## Supplementary Material

**FIG S1.**
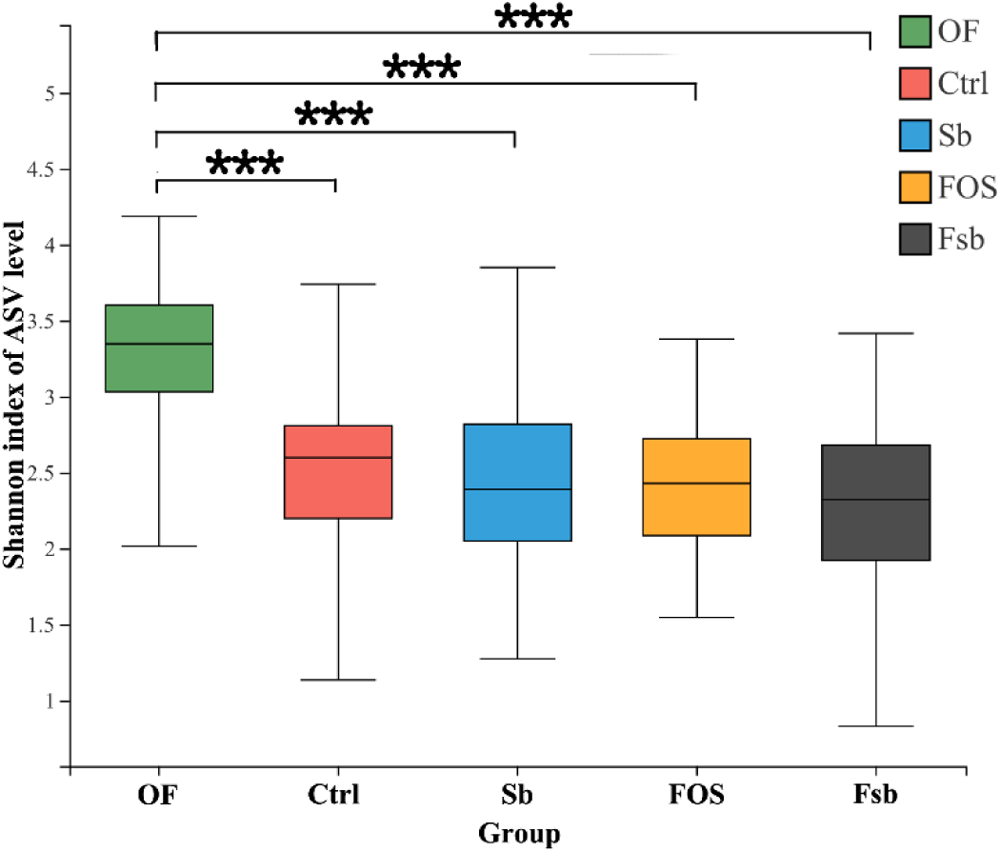
Shannon index of α-diversity. ANOVA, one-way analysis of variance. Statistical significance: ***, *P* ≤ 0.001.

**FIG S2.**
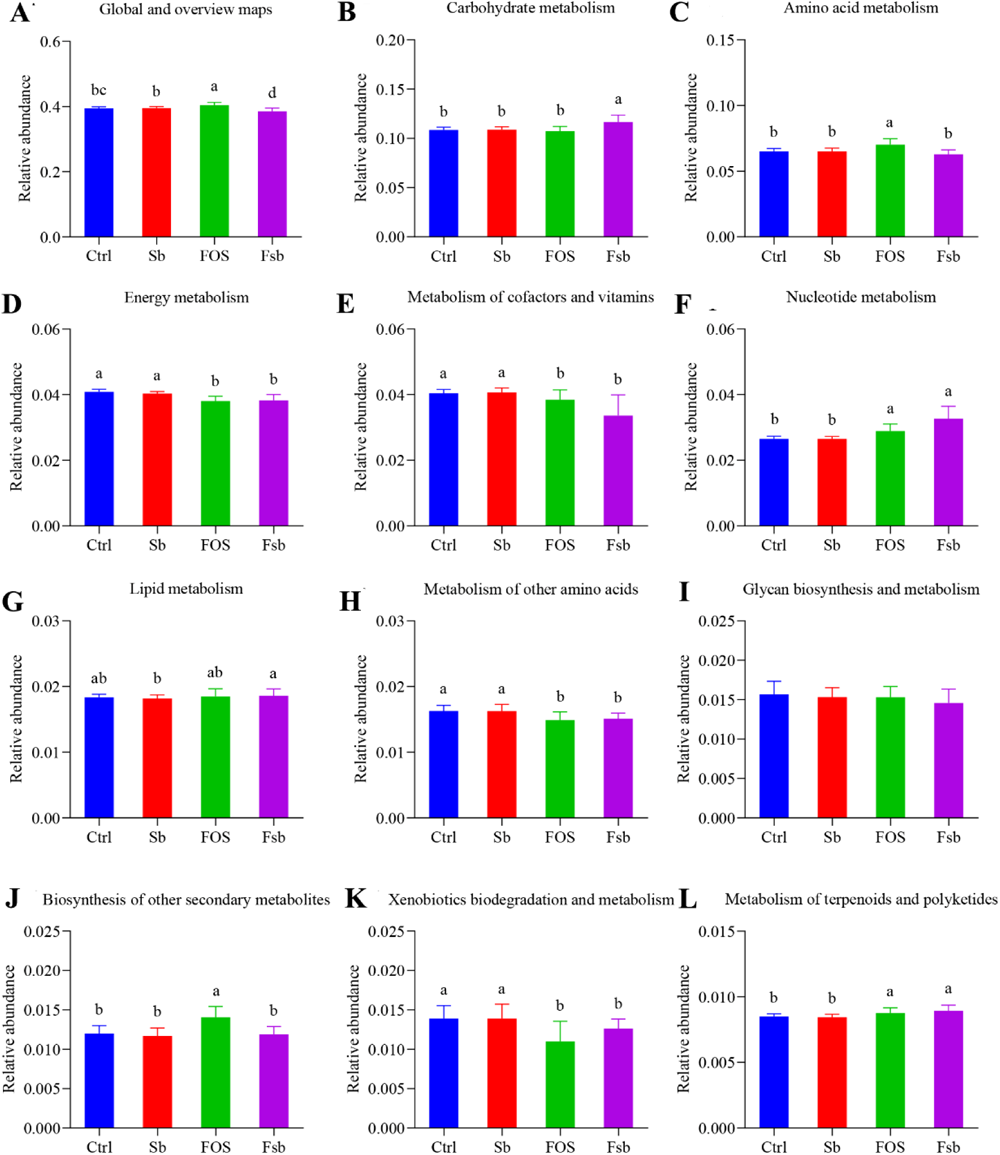
The differential KEGG functional pathways between all groups at Level 2. Different lowercase letters represent significant differences between different groups.

